# Diverse and novel lanthanide-binding PQQ-dependent enzymes

**DOI:** 10.64898/2026.07.10.737718

**Authors:** Marcos Y. Voutsinos, Colin M. Robinson, Rhys Grinter, Jillian F. Banfield

## Abstract

First described pyrroloquinoline quinone-dependent (PQQ) eight-bladed β-propeller proteins are calcium (Ca)-dependent, but many homologous bacterial enzymes are lanthanide (Ln)-dependent. Discovery of Ca-dependent six-bladed β-propeller PQQ-dependent dehydrogenases motivated the search for Ln-dependent six-bladed β-propeller PQQ-dependent enzymes in bacteria. Using *in silico* structural prediction of sequences from weathered rock, we identified ∼22,000 PQQ six-bladed β-propeller proteins in bacteria from 77 phyla, of which 63% of sequences have the active site residues needed to bind Ln. PQQ and La binding was biochemically confirmed for enzymes from uncultivated *Chloroflexi* and *Acidobacteria.* In structural models, Ln-binding periplasmic proteins interact with TonB-dependent transporters that may enable Ln uptake. Most genomes also encode predicted Ln-and PQQ-dependent eight-bladed dehydrogenases that clade with diverse alcohol and sugar dehydrogenases. Thus, Ln-dependent six and eight-bladed PQQ-dependent β-propeller proteins are implicated in diverse carbon substrate metabolisms in weathering rock and soil. We predict that many PQQ-dependent microbial enzymes are lanthanide dependent.

## Introduction

Pyrroloquinoline quinone (PQQ) dependent quinoproteins with six-and eight-blade β-propeller structures use PQQ, a quinone-containing prosthetic group, for electron transfer ^1^ ^2,3^. PQQ enzymes have been identified in organisms from all domains of life ^1,4,5^. They occur in many environments, including soils ^6–8^, seawater ^9,10^, the phyllosphere ^11^, and extreme environments ^12^. Eight-bladed PQQ calcium-dependent β-propeller proteins (PQQ-8β-Ca) oxidize a variety of substrates including methanol ^13^, ethanol ^14^, glucose ^3^, sorbose ^15^, antibiotics ^16^ and quinate ^17^. Eight-bladed PQQ lanthanide-dependent β-propeller enzymes (PQQ-8β-Ln) oxidize methanol ^12^, ethanol ^18^ and multi-carbon substrates (e.g., ExaF; ^18^, and PedH; ^19^. The active sites of PQQ-8β-Ln enzymes feature a lanthanide (Ln) binding motif containing two aspartate residues, the second of which does not occur in the Ca-dependent enzymes (MxaFI) ^20^; ^21^. Based on the presence of this motif, Ln-dependent methanol dehydrogenases were divided into five families (XoxF1-5) ^20^, with Ln coordination confirmed experimentally in XoxF1 ^22^, XoxF2 ^12^, XoxF4 and XoxF5 ^23^. Other predicted PQQ-8β-Ln alcohol dehydrogenases are divided into 11 families, but only Type 2b is confirmed to utilize lanthanides. Despite the fact that Ca concentrations far exceed Ln concentrations in soils (and Lns are sequestered in highly insoluble minerals ^24^), a predicted PQQ-8β-Ln methanol dehydrogenase was the most abundant protein detected in grassland soil ^6^. Overall, Ln dependency is generally predicted to be far more prevalent than Ca dependency in PQQ-8β proteins in natural environments ^10,20,25^. Preference for Ln-dependent relative to Ca-dependent methanol dehydrogenases has been attributed to increased enzyme efficiency ^12,26,27^. In fact, microbes capable of Ca-and Ln-dependent methanol oxidation (*mxaF, xoxF)* repress *mxaF* expression in favor of *xoxF* ^28^.

PQQ-dependent six-bladed β-propellers are calcium (Ca) dependent (PQQ-6β-Ca), and oxidize glucose ^3^, aldose ^29^; ^4^ and pyranose ^5^. Recently, the first eukaryotic PQQ-6β-Ca pyranose dehydrogenase was identified in the filamentous fungus *Coprinopsis cinerea* ^5^. It remains unclear whether PQQ-6β-Ln proteins exist, how they are distributed phylogenetically, and which environments they occur in.

Given the very low solubility of many Ln minerals, it is unsurprising that lanthanophores (siderophore-like molecules) are occasionally required for Ln acquisition. Uptake into cells can occur via TonB transporters ^30–35^. Lanthanides trafficked in the periplasm are expected to be tightly controlled to prevent precipitation and mismetalation ^36^. To date, three periplasmic Ln-binding proteins have been identified: lanmodulin ^37^, lanpepsy ^30^, and landiscerin ^38^. However, these are restricted to a small group of methylotrophs, and other proteins may be used by the much larger diversity of microorganisms with Ln-dependent enzymes.

In this study, we predicted the structures of PQQ-8β enzymes in weathered rock (and associated lichen) and soil, classified the structures as likely Ca-or Ln-dependent, and predicted likely substrate preferences. Motivated by the existence of PQQ-6β-Ca, and parallels involving PQQ-8β-Ca and PQQ-8β-Ln enzymes, we analyzed the active site residues of PQQ-6β proteins to search for evidence of PQQ-6β-Ln enzymes. Our analyses uncovered many putative PQQ-6β-Ln enzymes in soil and weathered rock. Thus, we searched public databases and identified many PQQ-6β-Ln enzymes in diverse bacteria and archaea. We expressed enzymes from Chloroflexi and Acidobacteria and confirmed Ln binding. Overall, the study extends Ln binding from PQQ-8β to PQQ-6β enzymes, indicates a wide diversity and distribution of PQQ-6β-Ln enzymes, and suggests that PQQ-dependent enzymes are often Ln-dependent.

## RESULTS AND DISCUSSION

### Bacteria in weathered granite encode diverse Ln-binding PQQ eight-bladed β-propeller proteins

We sampled a granite weathering profile to investigate the abundance and metal preference of PQQ-dependent enzymes. To provide context for studies of active site preference, we investigated Ca and Ln availability. Ca concentrations range from 22,000 ppm in slightly weathered granite to <5,000 ppm in associated soil. Total Ln concentration in all samples was <230 ppm (Table. S1, Data S1). Ca is predominantly in plagioclase and Lns occur in highly insoluble phosphate minerals such as rhabdophane ^8^. DNA was extracted from weathered rock, associated lichen and from soil and subjected to metagenomic sampling, sequence assembly, genome binning, and protein prediction. A total of 704 dereplicated genomes were reconstructed (Data S2).

We identified 15 clades of PQQ-8β-Ln. Notably, 485 PQQ-8β-glucose dehydrogenase genes were encoded by 175 genomes representing 10 phyla, and all contained the Ln-binding motif (Fig. 1, Fig. S1-3). Other PQQ-8β-Ln-dependent enzymes were classified as PQQ ADH Type 8 and XoxF3, both of which are implicated in oxidation of alcohols, including methanol. Methanol may be available from the atmosphere (e.g., methanol is the second most abundant organic gas in the troposphere; ^39^ and from aerobic oxidation of atmospheric methane and other VOCs ^40^ or from overlying soil, where it is produced by pectin breakdown.

**Figure 1.**
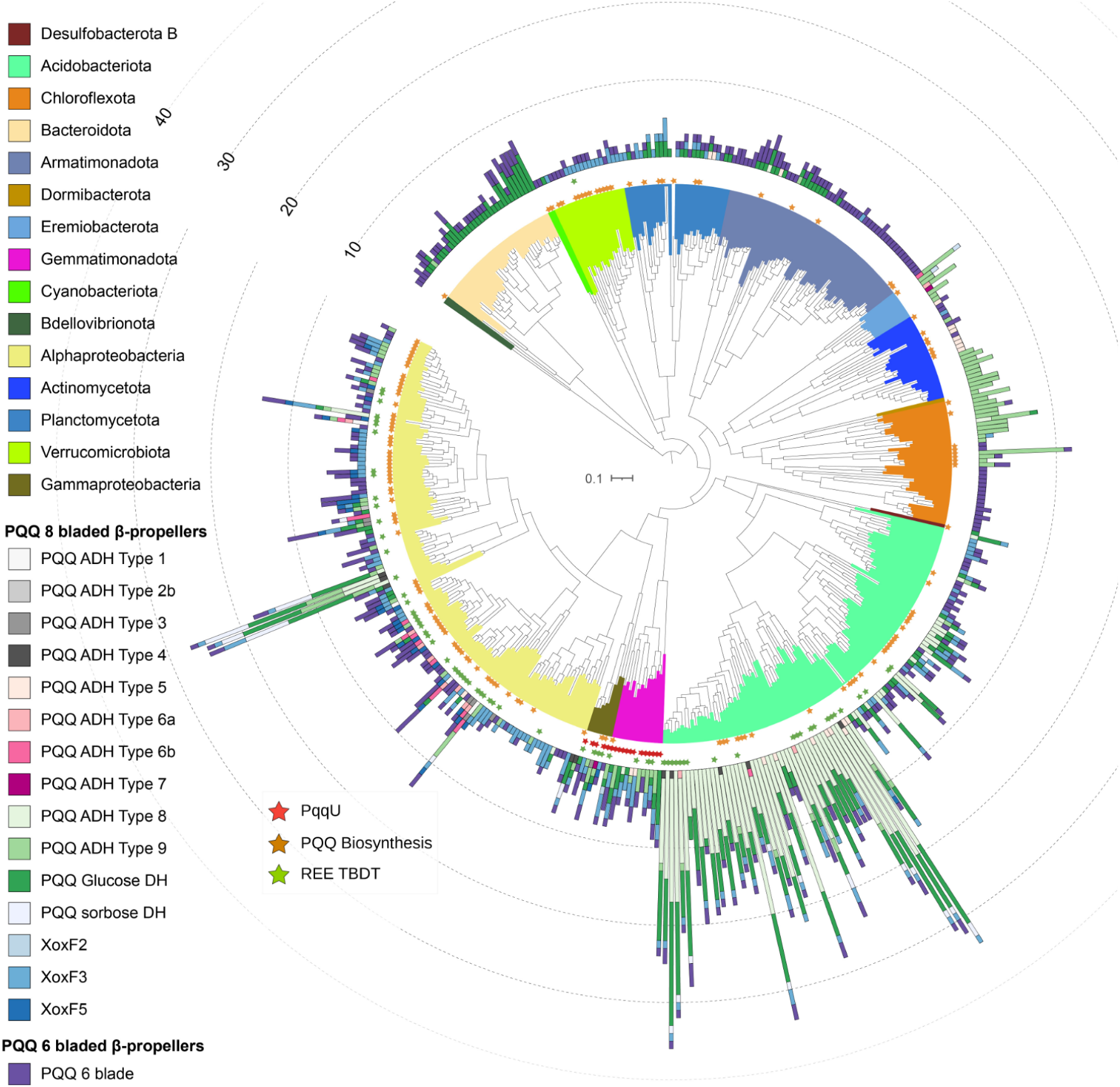
The majority of bacteria from weathered granite encode PQQ lanthanide dependent enzymes. Phylogenetic tree containing 472 genomes and their 2,107 PQQ enzymes (coloured bars). The concentric rings of stars indicate genomes that encode PQQ TBDTs (red stars), PQQ biosynthesis genes (orange stars) and Ln TBDTs (green star). For a rectangular view of the genome tree including full genome taxa see Figure S3.

We were curious as to the relative abundances of organisms that encode Ln-dependent enzymes in weathered rock microbiomes. We identified 4,935 non-dereplicated *rpS3* gene sequences from at least 23 phylum-level groups (Data S3). The highest PQQ dehydrogenase gene copy counts were found in PQQ-GDH, PQQ-ADH Type 8 and XoxF3, all of which belonged to the top 30% most abundant species (Fig. S4). These species belonged to common soil phyla, including Gemmatimonadota, Armatimonadetes, Alphaproteobacteria, Actinomycetes, Chloroflexi, and Acidobacteria.

### Putative novel periplasmic lanthanide cargo proteins interact with TonB transporters

Protein clustering methods have shown that TonB-dependent transporters (TBDTs) primarily form groups based on substrate activity ^41^. Using a sequence similarity network analysis of all 7,472 binned TBDTs, we identified 74 TBDTs that clustered closely with five TBDTs (Fig. 2A) previously shown to import Ln (TBDT-Ln) without transporting PQQ ^32,34,35,42^. These 74 Ln-TBDT sequences were misannotated as YnCD (now PqqU) and the top PDB hit was PqqU (PDB: 6V81) but they did not contain the conserved PQQ binding residues ^43^ (Data S4). We did not identify any TBDT-Ln that clusters with the TBDT that is expected to take up methylolanthanin (*mluA*) ^44^. The 74 TBDT-Ln were encoded in the genomes of Alphaproteobacteria, Gammaproteobacteria, Acidobacteria, Gemmatimonadetes, and Verrucomicrobia (Fig. 1). Analysis of the TBDT-Ln genomic context revealed neighbouring PQQ biosynthesis genes and a small hypothetical protein (∼120 aa, ∼8 kDa) (Fig. 2B) almost always encoded next to Ln-TBDT. The hypothetical protein has a Sec/SPI signal peptide, indicating transport to the periplasm. It does not contain any conserved domains and the modelled structure has no closely related structures in PDB. Using Chai-1, the protein is predicted to bind La^3+^ in a region containing three aspartate, one glutamate and one threonine residue, potentially allowing for coordination with eight oxygens, highly similar to Ln binding in lanmodulin (Fig. 2C) and XoxF. Analysis of models for six examples revealed strong interaction (avg. buried surface area = 1709 Å²) between the small proteins and the TBDT periplasmic cavity (Fig. 2D, Fig. S5). Sometimes the protein was encoded separately, but still showed strong interaction with TBDT-Ln and in some cases the protein did not bind to the periplasmic cavity unless La^3+^ was present (RPR_22_05_09_5_L_scaffold_393_57). One model showed good interaction confidence between the small protein and the plug and barrel of the TBDT and complimentary electrostatic surfaces (RPR_22_05_09_4_k127_6384731_3) however, despite multiple models showing similar interaction geometry they showed generally poor ipTM values and interaction confidences (Data S4). TBDTs often function in synchrony with extracellular and periplasmic substrate binding proteins ^45^. Based on our findings we predict that TDBT-Ln imports Ln and passes it to the small periplasmic Ln-binding cargo protein, which likely then transfers it to a quinoprotein. Similar interactions occur during import of cobalamin (Vitamin B_12_) and iron via TBDT ^46^ ^47^.

**Figure 2.**
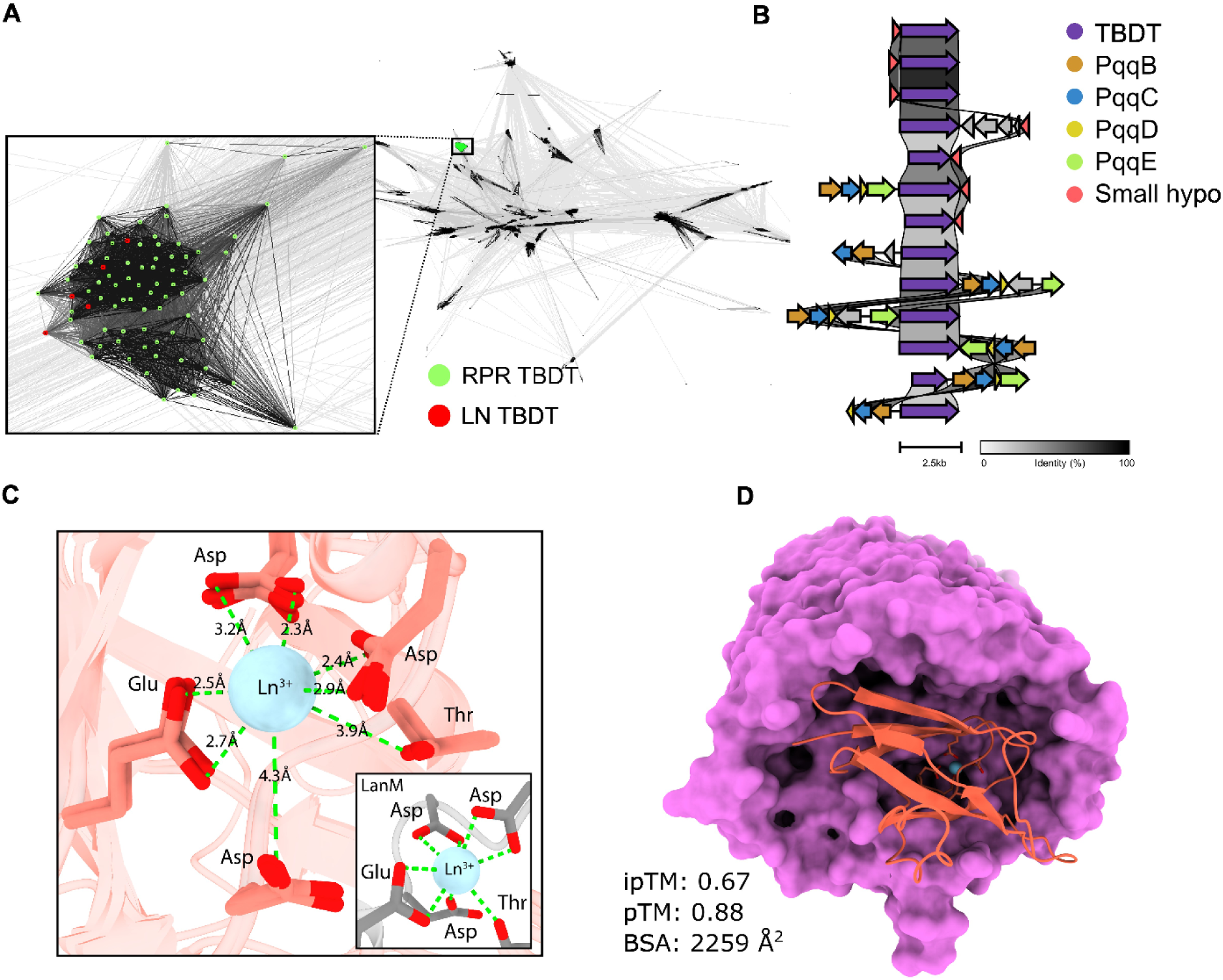
A small hypothetical periplasmic protein is predicted to bind lanthanides and interact with a putative lanthanide TonB dependent transporter (Ln-TDBT). **A)** Sequence similarity network of more than 7,000 TBDT homologs derived from 704 genomes clustered with five experimentally confirmed Ln-TBDT reference sequences. Each point represents an individual protein, grey lines represent BLASTp e-values > 1×10^-10^, with increasing line darkness indicating increasing sequence similarity. The insert shows 74 TBDTs (RPR TBDT, green) from genomes clustering with Ln-TBDT (red). **B)** Diagram showing loci with RPR TBDTs and adjacent PQQ biosynthesis operons and/or a small hypothetical periplasmic protein **C)** Predicted interaction of four superimposed small hypothetical periplasmic proteins with Ln^3+^. Insert displays lanthanide interaction of lanmodulin, PDB: 6MI5. **D)** Predicted interaction between an RPR TDBT from Alphaproteobacteria (RPR_22_05_09_4_k127_6384731_2, lavender), the adjacent encoded small periplasmic protein (RPR_22_05_09_4_k127_6384731_3, salmon) and Ln^3+^ (blue sphere).

### Ln-dependent PQQ six-bladed β-propeller dehydrogenases

We used the nine PQQ binding residues (Arg^273^, Hist^285^, Arg^430^, Asn^431^, Glu^539^, Glu^560^, Trp^563^, Asn^564^, Arg^622^) of *Coprinopsis cinerea* PQQ-6β pyranose dehydrogenase (CcPDH, PDB: 6JWF) as a signature of PQQ dependency in 6-bladed β-propeller proteins ^5^. In the genomes isolated from weathered granite we identified 2,290 six-bladed β-propeller proteins, 483 of which have all nine expected PQQ binding residues (Fig. 1 & Fig. 3A). These 483 sequences derived from 11 phyla: Alphaproteobacteria, Gammaproteobacteria, Cyanobacteria, Armartimonadetes, Gemmatimonadetes, Bdellovibrio, Verrucomicrobia, Planctomycetes, Actinobacteria, Acidobacteria and Chloroflexi (Fig. 3B). The PQQ binding residues are predicted to be positioned with the correct geometry for PQQ coordination (Fig. 3C-D) (pLDDT > 0.9, Data S5), thus we conclude that these proteins are PQQ dependent. Predicted structures are most similar in structure to CcPDH (RMSD between 215 pruned atom pairs was 1.02 Å, 62% of sequence) (Fig. 3E). Superimposition of active sites revealed that, unlike the Ca^2+^ coordinating residues (Asn^448^, Ser^449^, Leu^450^, Asp^451^, Glu^471^) of CcPDH (Fig. 3F), ∼80% of the bacterial homologs have Asn, Glu, Arg, Asp, and Asp (Fig. 3F). Modelling placed La^3+^ in the active site (Fig. 3F) with high confidence (pLDDT > 0.9, Data S5). The Ca*^2+^*in CcPDH coordinates with six oxygens of which two are from water molecules (Table S2) ^5^. Lanthanides are coordinated by 7-9 oxygens in PQQ-8β ^48^. The metal binding active site of bacterial PQQ-6β appears to support coordination with at least 7 oxygen atoms (Table S2) and more may be provided by water molecules. All sequences contained an N-terminal Sec/SPI signal peptide domain indicating export of the protein from the cytoplasm. Many putative PQQ-6β Ln-dependent enzymes contained Auxiliary Activity Family 12 (AA12) domains, indicating CAZyme activity (Data S6), as identified in CcPDH ^5^. To our knowledge, bacterial PQQ-6β with AA12 domains have not been previously reported ^49^. Many of these proteins were predicted to be NHL-repeat protein and L-sorbose dehydrogenases (both are beta propeller proteins).

**Figure 3.**
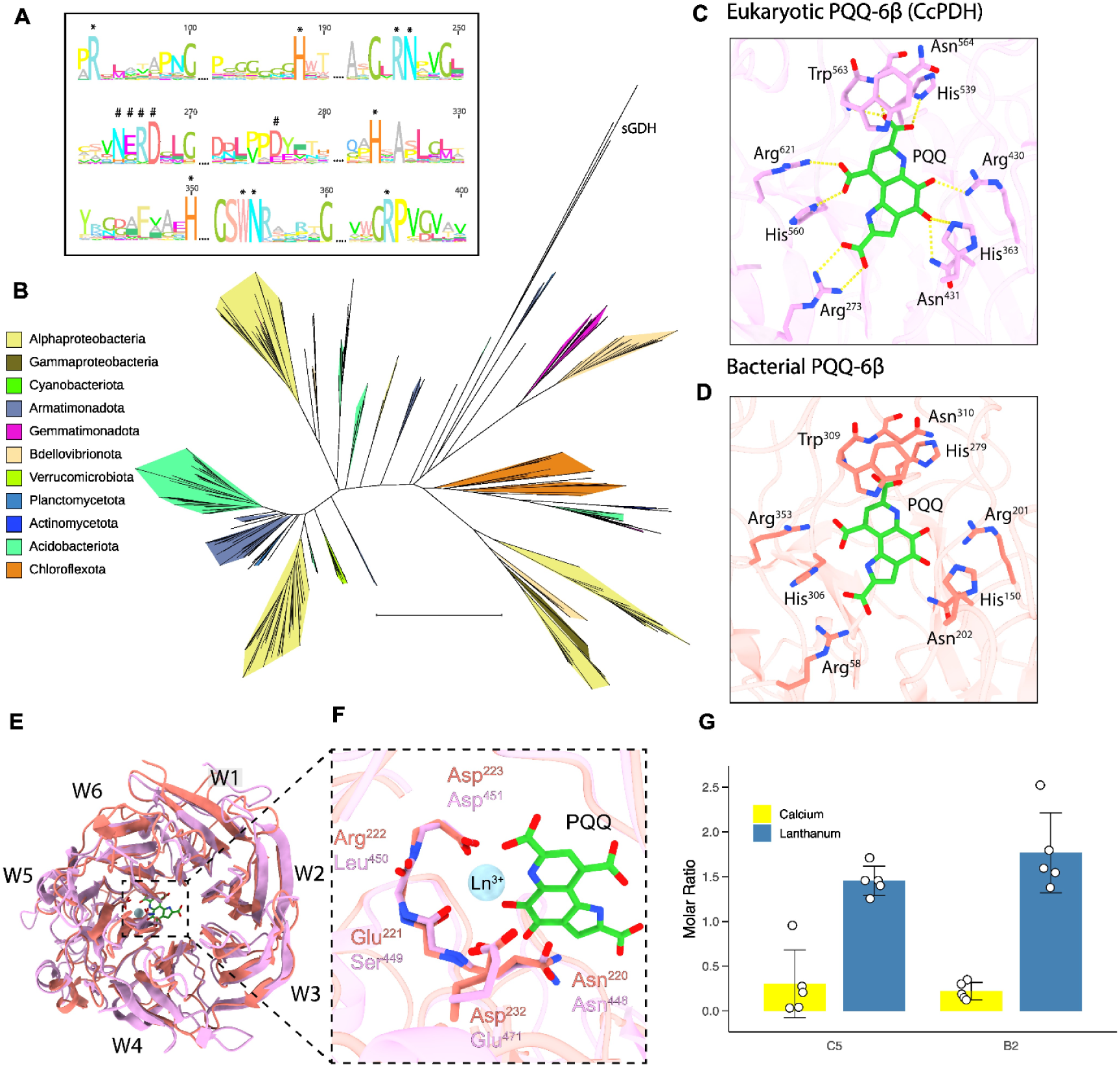
Bacterial six-bladed β-propeller enzymes from weathered granite are PQQ and lanthanide binding. **A)** Consensus sequence logo of all 378 sequences highlighting the conserved residues responsible for binding PQQ (‘*’) and a metal cofactor (‘#’). **B)** Phylogenetic tree of 378 binned PQQ six-bladed β-propeller sequences and 3 soluble six-bladed PQQ glucose dehydrogenase reference sequences (labelled ‘sGDH’, PDB IDs: 1C9U, 8RG1 and 1QBI). **C)** The nine residues involved in PQQ binding of Eukaryotic CcPDH (PDB: 6JWF) **D)** Predicted residues of a PQQ six-bladed β-propeller protein, RPR_22_05_09_1_scaffold_1078_6 (C5) from an Acidobacteria genome, involved in PQQ binding. **E)** C5 modelled with PQQ and La^3+^ and superimposed with its foldseek best hit, CcPDH (PDB ID: 6JWF; E-value = 8.03e-25; RMSD = 0.995 Å out of 212/343 atom pairs). The six antiparallel β-sheets are labelled W1-W6. **F)** Zoomed-in view of the active site illustrating the residues involved in lanthanum (blue sphere) coordination. Only the backbone carbonyl oxygen of Glu221 and Arg222 is shown and predicted to be involved in metal coordination. **G)** Molar ratio of metals in C5 and B2 proteins after incubation with calcium (yellow) and lanthanum (blue). White circles are the molar ratio of the five technical replicates, bar plots are the mean and the error bars are the standard deviation.

### Biochemical confirmation of La binding by PQQ-6β proteins

To confirm PQQ binding, two predicted PQQ-6β-Ln proteins from phylogenetically defined clades were produced in *E. coli*, purified, reconstituted with PQQ, La^3+^, and Ca^2+^, and subjected to ultraviolet-visible light spectroscopy (UV-Vis) and inductively coupled plasma-mass spectroscopy (ICP-MS). Protein B2 from a Chloroflexi order Thermomicrobiales genome (RPR_22_05_09_4_k127_6634144_13) eluted at a retention volume of ∼130 mL (Fig. S6A) and was ∼55 kDa (expected weight ∼49 kDa) (Fig. S6B) and protein C5 from Acidobacteria family Bryobacteraceae genome (RPR_22_05_09_1_scaffold_1078_6) eluted at a retention volume of ∼40 mL (Fig. S6C) and was ∼49 kDa (expected weight ∼ 47 kDa) (Fig. S6D). The identity of both proteins was confirmed by liquid chromatography-mass spectrometry. Under UV-Vis illumination, the spectrum from apo-B2 and apo-C5 displayed a peak at ∼280 nm. When apo-B2 and apo-C5 were incubated with PQQ_aq_ the UV-Vis spectrum displayed peaks at ∼330 nm and ∼380 nm (Fig. S6E). These peaks are broadly consistent with those reported for both six-^4,29^ and eight-bladed ^12,50^ quinoproteins when bound to PQQ, which generally display one peak between ∼330 and 360 nm and sometimes a secondary peak at ∼375-400 nm. The peaks at 330 nm from B2 and C5 are exactly comparable to the peak at ∼330 nm reported for PQQ-bound to the 6β-Ca enzyme CcPDH ^51^. We conclude that the nine PQQ binding residues were indeed a signature of PQQ binding in bacterial PQQ-6β proteins.

To test for La binding and preference for La^3+^ over Ca^2+^, we co-incubated the B2 and C5 PQQ enzymes with Ca^2+^ and La^3+^. The ICP-MS results revealed a preference for La over Ca (Fig. 3G), supporting the metal binding active site NERD…D and NGRD…D motifs as a signature of Ln binding. Specifically, protein C5 exhibited ∼1.5 La and ∼0.25 Ca atoms per monomer and protein B2 exhibited ∼1.8 La and ∼0.25 Ca atoms per monomer. A small degree of Ca^2+^ binding is also observed in other Ln-dependent PQQ enzymes ^12^. As the total metal atom content per monomer is >> 1, we infer some non-specific binding or the presence of a second metal binding site. In structural models of B2 predicted with one La and one Ca per monomer, Ca was incorporated into a second metal binding site between two of the six B-sheets. The predicted metal binding residues were a commonly occurring motif (DxD/NxDxxxD/E) across the 21,992 PQQ-6β multiple alignment (Fig. S7). This second metal site may serve a structural role, possibly stabilising the propeller structure, as is observed in other six-bladed PQQ β-propeller structures ^4,29^.

### Predicted biological context of PQQ-6β-Ln proteins

Commonly a monoheme cytochrome is encoded adjacent to the PQQ-8β gene (e.g., cytochromes ^C^L known as MxaG and XoxG involved in methanol oxidation ^52^. In Alphaproteobacteria and Gammaproteobacteria, PQQ-6β-Ln proteins were commonly adjacent to diheme cytochrome c class I genes (cytochrome C4). The cytochrome C4 sequences contain a Sec/SPI signal peptide indicating transport to the periplasm. Chai-1 was used to model the cytochrome with two heme C groups, two Fe^2+^, the 6β protein, PQQ, and La^3+^. The cytochrome and PQQ-6β-Ln protein are predicted to interact with high confidence (iPTM 0.93), with a buried surface area of 1270 Å² (Fig. 4A-C, Data S7). The cytochrome docks on top of the 6β protein adjacent to the active site, such that the PQQ and heme c groups are ∼12.5 Å apart. Based on the PQQ and heme distances in PQQ quinohemoprotein ADH-IIG (PDB: 1YIQ, ^53^, PQQ and heme C are close enough to enable electron transfer (∼12 Å). The Alphaproteobacteria cytochrome C4 shares a high structural similarity (RMSD of 0.809 Å between 140 pruned atom pairs) to cytC4 in the cytochrome bc1-cbb3 supercomplex (PDB: 8SMR) from *Pseudomonas aeruginosa* ^54^.

**Figure 4.**
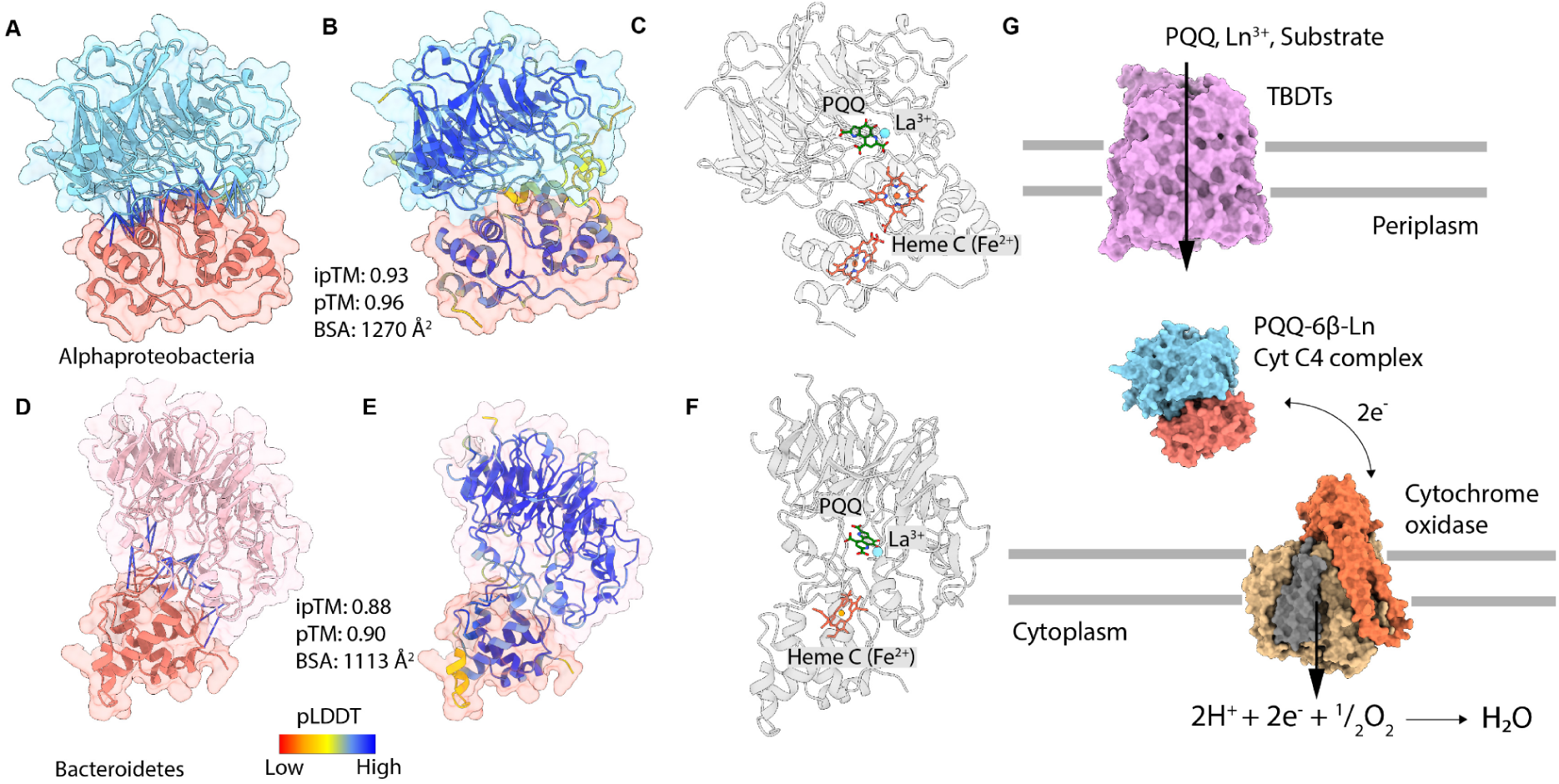
Interactions between PQQ-6β-Ln and the di-heme Cytochrome C4 and mono-heme Cytochrome C_551_. **A)** Alphaproteobacterial PQQ-6β-Ln (blue) and Cytochrome C4 (salmon) interact with high confidence **B)** The ribbons of PQQ-6β-Ln (blue surface) and Cytochrome C4 (salmon surface) are coloured according to their predicted local distance difference test (pLDDT). **C)** Binding sites of PQQ, La^3+^, two heme C and two Fe^2+^ in the complex. PQQ-6β-Ln and Cytochrome C4 and greyed out. **D)** Bacteroidete PQQ-6β-Ln (pink) and Cytochrome C_551_ (salmon) interact with high confidence **E)** The ribbons of PQQ-6β-Ln (pink surface) and Cytochrome C4 (salmon surface) are coloured by pLDDT **F)** Binding sites of PQQ, La^3+^, heme C and Fe^2+^ in the complex. PQQ-6β-Ln and Cytochrome C_551_ and greyed out. **G)** Simplified model of predicted electron transfer from the periplasmic PQQ-6β-Ln Cytochrome C4 complex to a membrane bound terminal cytochrome oxidase.

Across multiple orders of Bacteroidetes including Sphingobacteriales, Chitinophagales, and Cytophagales, PQQ-6β-Ln were commonly adjacent to periplasmic monoheme cytochrome c_551_ (Cytc_551_). Chai-1 was used to model the cytochrome with a single heme C, Fe^2+^, the 6β protein, PQQ, and La^3+^. The cytochrome and PQQ-6β protein are predicted to interact with high confidence (iPTM 0.88, Fig. 4D-F, Data S7), with a buried surface area of 1113 Å². The Bacteroidetes cytochrome c551 shares a high structural similarity (RMSD of 0.898 Å between 89 pruned atom pairs, 70% of Bacteroidetes Cytc_551_ sequence) to the cytochrome c551 fused to copper nitrite reductase (PDB: 3ZBM) from Betaproteobacteria *Ralstonia pickettii* ^55^.

To our knowledge, interaction of a PQQ quinoprotein with a cytochrome C4 has not been previously reported. Results suggest that the Alphaproteobacteria and Bacteroidetes PQQ-6β enzymes are involved in an electron transfer complex within a respiratory pathway (Fig. 4G). Interestingly, some bacterial PQQ-6β proteins contain a heme-binding domain that may serve essentially the same purpose. This apparent fusion is reminiscent of the heme-binding domains that occur in some 8-bladed β-propeller proteins (e.g., Type IV alcohol dehydrogenase quinohemoproteins).

### PQQ-6β proteins are encoded by bacteria from 77 phyla and cluster by active site metal cofactor

To determine how widespread PQQ-6β-Ln proteins are in bacteria, we searched for them in the 107,235 bacterial species clusters of the Genome Taxonomy Database (GTDB). We used the nine PQQ binding residues of 6JWF as a signature of PQQ dependency and the presence of NERD…D of C5 and NGRD…D of B2 in the active site as a signature of Ln binding.

In total 11,203 bacterial genomes encoding 21,992 PQQ-6β sequences spanning 77 phyla were identified (Fig. 5A, Data S8) of which the most common metal binding motifs were NERD…D (49%) and NGRD…D (13%) (Fig. 5B) spanning 72 phyla, suggesting that the majority of identified PQQ-6β proteins are Ln-dependent. Many of these phyla have not been previously associated with PQQ-dependent enzymes or Ln binding, highlighting that PQQ-6β-Ln are widespread.

**Figure 5.**
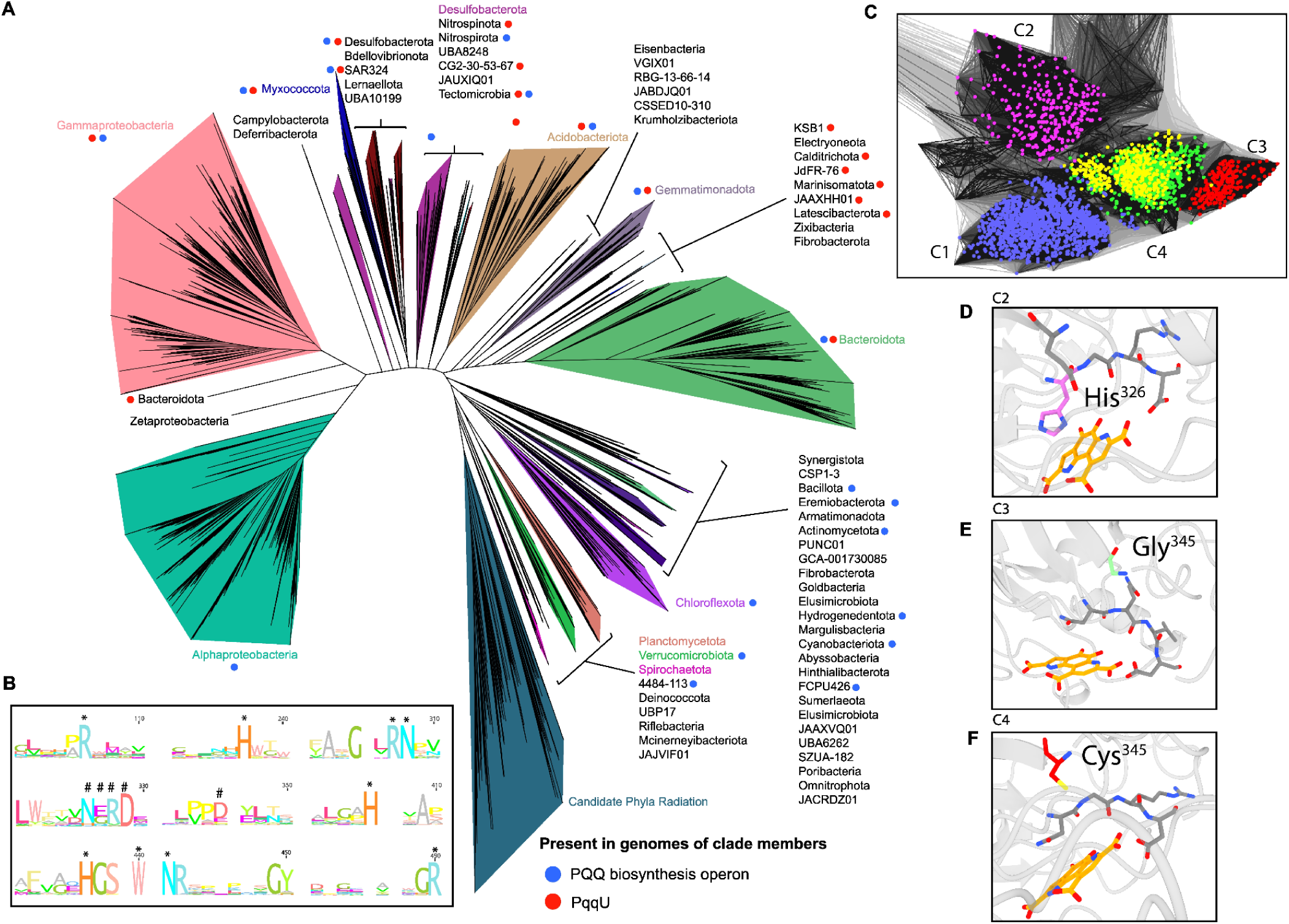
Bacterial PQQ-6β enzymes are encoded by bacteria from 77 phyla and mostly cluster according to their metal dependence. A) 16,726 genomes harbour 21,992 PQQ 6-bladed β-propeller enzymes. B) Consensus multiple alignment of all 21,992 sequences, PQQ binding residues (asterisk) and metal cofactor binding residues (hash symbol) are indicated. C) Sequence similarity network of 3,732 sequences (one per genus) decorated by the most common metal binding motifs; NERD…D (C1), HGRD…E (C2), NQCD…G (C3), NGRD…D and NGRD…C (C4). D), E) and F), Metal binding active sites of representative sequence from C2, C3 and C4 clusters in sequence network, respectively.

We were curious to know whether metal preference is predictable based on sequence similarity. This is the case for some PQQ-8β enzymes (e.g., XoxF vs MxaF and PedH vs PedE; ^19,20^, but not for other phylogenetically defined PQQ-8β enzyme clades (e.g., Types 2b and 6a alcohol dehydrogenases) that are predicted to include both Ca and Ln-dependent proteins ^56^. In a PQQ-6β protein sequence similarity network generated using 3,732 sequences (one per genus), 88% of sequences formed four distinct clusters (Fig 5C). The sequences within each cluster derive from many different phyla, suggesting that all principal PQQ-6β variants are widely dispersed in Domain Bacteria.

In cluster 1 (C1, Fig. 5C), 93% of the sequences contain N326, E327, R328, D329, D345 (98% of all NERD…D containing sequences) and these proteins likely bind Ln. In cluster 2 (C2, Fig. 5C), 97% of the sequences contain H326, G327, R328, D329, E345 (99% of all HGRD…E containing sequences). Active site modelling indicates H326 may change the metal coordination geometry and reduce the number of available coordinating oxygens (Fig. 5D). Thus, we infer that Cluster 2 proteins do not bind Ln. In cluster 3 (C3, Fig. 5C), 86% of sequences contain N326, Q327, V328, D329, G345 (100% of all NQVD…G containing sequences). Modelling of the active site indicates that the Gly^345^ residue may not be involved in Ln coordination because the backbone carbonyl group is distant from the metal cofactor (∼10 Å) (Fig. 5E).

In cluster 4 proteins, 57% of the sequences have the active site residues N326, G327, R328, D329, D345, and we infer that they are Ln binding. 24% of cluster 4 sequences contain N326, G327, R328, D329, C345. Chai-1 modelling of the NGRD…C active site indicates that the sulfur group of Cys^345^ in Cluster 4 proteins has the required geometry for interaction with a metal group (Fig. 2F). However, lanthanide binding is unlikely because lanthanides are hard Lewis acids that favour oxygen donors, whereas cysteine is a soft Lewis base due to its thiol (-SH) group.

The network topology is based on the entire protein sequence, thus likely groups together proteins with shared evolutionary history. The network findings suggest that, for proteins that place in cluster 1, Ln binding is often predictable based on the overall protein sequence. Apparently, over time, switching between Ln and dependence on another metal does not often occur in clusters 1, 2 and 3 proteins. The admixture in Cluster 4 suggests a relatively facile transition from Ln to another metal.

### Some PQQ-6β-Ln proteins are associated with predatory bacteria and obligate symbionts

PQQ-6β-Ln proteins were encoded in two newly reconstructed high-quality MAGs for Bdellovibrionota (Fig. 1) and in many public *Bacteriovorax* and *Bdellovibrio* genomes (Fig. 5A), including representatives experimentally shown to have obligate predatory, intracellular lifestyles ^57–59^. Recent work shows Ln amended soils increased the relative abundance of *Bdellovibrio* relative to the non-Ln amended soils ^60^. The proteins were most similar to an extracellular PQQ-6β-Ca CAZyme from fungus, *Trichoderma reesei* (*Tr*AA12, PDB ID: 6I1T) ^61^ that preferentially oxidises L-fucose ^61^ found in extracellular polysaccharides and outer membrane lipopolysaccharides of certain Gram-negative bacteria ^62^. It is possible that the PQQ-6β-Ln proteins are involved in oxidation of L-fucose associated with the surfaces of bacterial prey.

We identified 247 genomes representing bacteria of 12 classes of the Candidate Phyla Radiation (CPR, sometimes referred to as Patescibacteria) that encode PQQ-6β-Ln proteins. All contain one or two pairs of cysteine residues that are predicted to form disulphide bonds providing additional stability. No CPR bacterial genomes encode any PQQ biosynthesis genes (Fig. 5A). Bacteria biosynthesize PQQ via the PQQ operon ^63^, but due to its small size (330 Da), PQQ can be lost via diffusion from the cell. *Escherichia coli* ^64^ and many other Gram-negative bacteria do not biosynthesise PQQ but instead import PQQ via the TonB dependent transporter, PqqU ^43^. CPR bacteria generally have ultra-small cells, extremely streamlined genomes (∼1 Mbp) and episymbiotic lifestyles, with extensive metabolic dependence on their host bacteria ^65,66^. They are predicted to lack an outer membrane (i.e., they are not diderm) and thus outer membrane transport systems such as TonB dependent transporters. Given evidence for direct cytoplasmic contact between (at least some) CPR and host cells ^67^, they likely derive PQQ directly from their hosts.

### Archaea encode six-bladed β-propellers with conserved PQQ binding residues

We identified 153 different PQQ-6β proteins in genomes from Halobacteriota, Nanoarchaeota, Aenigmatarchaeota and one genome from each of Micrarchaeota and Thermoplasmatota (Data S9). The consensus metal binding active site residues (Asn, Glu, Arg, Asp, Asp) indicate Ln dependency (Fig. S8A, Data S9). All nine predicted PQQ binding residues in all of the archaeal proteins are shared with PQQ-6β pyranose dehydrogenase from *Coprinopsis cinerea* (Fig. S8B), and they are completely conserved in related PQQ-dependent enzymes from Bacteria and Eukaryotes. The Ln-binding motif shares the same geometry as the Ln binding Bacterial homologs (Fig. S8C). An unrelated PQQ-dependent aldose sugar dehydrogenase (PDB: 3A9H) predicted to bind Ca in the metal binding site was previously identified in the hyperthermophilic archaeon *Pyrobaculum aerophilum* ^4^. This enzyme shares structural similarity and identical PQQ binding residues with soluble PQQ aldose sugar dehydrogenase from *E. coli* (PDB: 2G8S), except for an additional arginine not present in the *E. coli* enzyme. The newly described archaeal PQQ-6β enzymes share only 27% sequence identity with the previously described archaeal enzyme. Given this, and PQQ binding residues that are divergent from those PQQ aldose sugar dehydrogenase, we suggest their classification as a new class of putative Ln-dependent PQQ-6β quinoproteins.

### Association of PQQ-6β-Ln proteins with loci involved in carbohydrate utilization

To gain insight into the function of the PQQ-6β-Ln proteins we screened their genetic organization across 3,732 genomes. We found 400 genomes encoding PQQ-6β within CAZyme gene clusters (CGCs) across 30 bacterial phyla (Fig. S9, Data S6) and Halobacteriota and Aenigmattarchaeota. The CGCs contained CAZymes, peptidases, sulfatases, transporters, transcription factors and signal transduction proteins commonly involved in carbohydrate utilisation ^68^. We also identified five examples of PQQ-6β CGCs sharing similarity with biochemically characterised polysaccharide utilisation loci known to breakdown xylan, arabinogalactan, beta-glucan, pectin, and arabinan. (Fig. 6).

**Figure 6.**
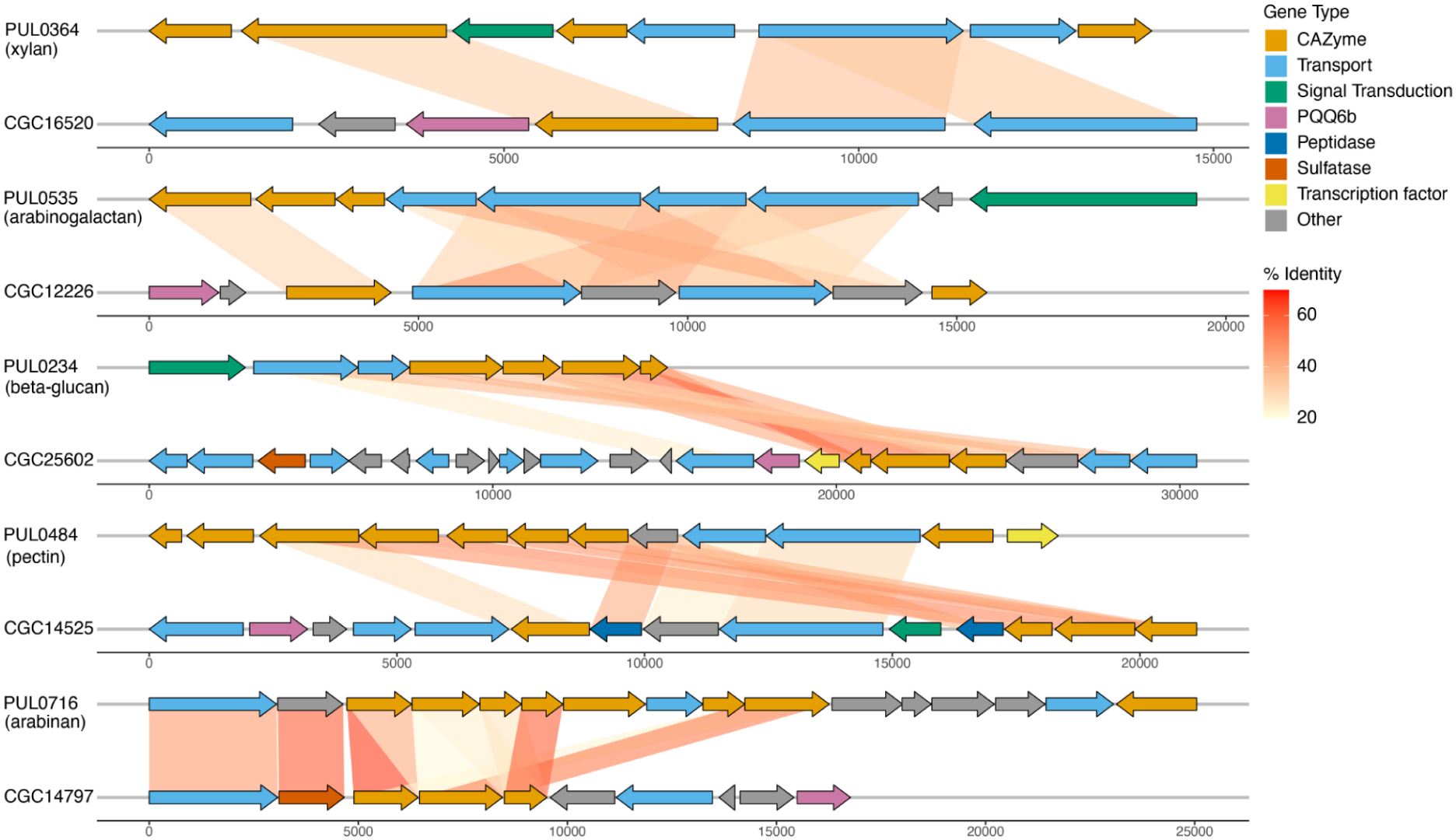
Predicted substrates of PQQ-6β proteins. Gene cluster comparisons involving sequences from this study (CGC) and biochemically characterised polysaccharide utilisation loci are shown pairwise, with inferred protein functions and amino acid identity indicated for homologs. Gene scale is in base pairs.

The analogous eukaryotic PQQ-6β-Ca enzyme CcPDH is believed to be involved in extracellular carbohydrate degradation. We speculate that membrane bound polysaccharides are processed and oligosaccharides are transported into the periplasm where reducing-end hydroxyl groups are oxidized by PQQ-6β and electrons shuttled to cytochromes ultimately for utilization in the TCA cycle. Given the genomic context, we suspect that these genomic loci are involved in uptake of soil-associated carbon compounds that are then oxidized by PQQ-6β proteins. Overall, the carbohydrate degrading function of CcPDH and association of the newly reported bacterial PQQ-6β-Ln genes with genes encoding carbohydrate utilisation and transport proteins suggests general functions of PQQ-6β-Ln proteins in carbohydrate metabolism.

## Conclusion

Here, we report previously unknown PQQ-6β-Ln proteins in diverse bacteria and some archaea and confirmed PQQ binding and Ln specificity for two of these proteins experimentally. Notably, this suggests that PQQ residues are conserved in all three domains of life and Ln-binding residues are conserved in bacteria and archaea. The experimental confirmation of *in silico* structure-based active site predictions supports the wider use of active site residues to predict metal binding preference. We deduce that metal preference in three of four clusters of enzymes in sequence similarity based networks can be predicted based on cluster assignment. Genomic context analyses extend the finding of carbohydrate degradation function of eukaryotic PQQ-6β-Ca proteins to bacterial and archaeal PQQ-6β-Ln proteins. Both heme-binding motifs in some proteins and multimer predictions that include mono/diheme cytochromes encoded adjacent to some PQQ-6β-Ln proteins suggest that electrons pass from the substrate to PQQ and then via a cytochrome to a terminal oxidase. In weathering granite and granitic soil, PQQ-6β and PQQ-8β proteins appear to be prevalent and primarily lanthanide dependent. In fact, overall, we suggest that most PQQ-dependent proteins are Ln-dependent.

## Methods

### Sample locations and sample collection

We sampled progressively weathered I-type granodiorite that outcrops in central Victoria, Australia (-37.3220, 142.8296). The densities of weathered and fresh rocks were measured and compared to provide an indication of the degree of weathering. Five samples were collected in triplicate. Sample 1 was moderately to highly weathered (2.1 g/cm^3^) and collected from the base of the weathered granite profile, about 1-3 cm below the soil zone. Sample 2 was moderately weathered (2.2 g/cm^3^) and 3-5 cm above sample 1. Sample 3 was slightly to moderately weathered (2.3 g/cm^3^) and 5-8 cm above sample 2. Sample 4 was slightly weathered (2.4 g/cm^3^) and collected from the top of the granite boulder about 5-8 cm above sample 3. Sample 5 was a lichen colony (thallus) collected from the surface of weathered granite 3-5 cm above sample 4.

### Whole-rock chemical analysis

Whole rock chemistry of the samples were determined using inductively coupled mass spectroscopy (ICP-MS) (Data S1), as previously described in Voutsinos et al ^8^.

### DNA extraction, sequencing and metagenomic assembly

Samples from the five weathered regions were collected for biological analysis in triplicate into 50 ml centrifuge tubes and flash frozen in liquid nitrogen before being transported to the lab on dry ice and stored at-80 degree celsius on the same day. DNA was extracted from 10 grams of sample using the PowerMax Soil DNA isolation kit (Qiagen). Metagenomic library preparation and DNA sequencing were performed at AGRF, Melbourne on a NovaSeq 6000 platform, producing 250 bp paired-end reads with a target inter-read spacing of 550 bp. Raw reads were trimmed of Illumina adapters and PhiX and other Illumina trace contaminants were removed using BBTools (https://github.com/bbushnell/BBTools) with default settings. Reads were quality trimmed using Sickle (https://github.com/najoshi/sickle) with the following parameters:: pe-q 20-l 20. Samples were co-assembled using IDBA-UD ^69^ with default settings and the --pre_correction parameter. Where IDBA-UD would hit a memory wall due to high sample complexity, MEGAHIT ^70^ was used with default settings and the following parameters: --presets meta-large --min-contig-len 1000.

### Metagenome annotation

Pullseq was used to filter samples and remove contigs smaller than 1000 bp (https://github.com/bcthomas/pullseq). Open reading frames (ORF) were predicted on all contigs using Prodigal v2.6.3 ^71^ with the following parameters:-m-p meta. Predicted ORFs were annotated using USEARCH ^72^ to search all ORFs against Uniprot ^73^, Uniref90 and KEGG ^74^. 16S ribosomal rRNA genes were predicted using the 16SfromHMM.py script from the ctbBio python package using default parameters (https://github.com/christophertbrown/bioscripts) and the ssu-align-0p1.1.cm database. Transfer RNAs were predicted using tRNAscan-SE. The metagenomes and their annotations were then uploaded to ggkbase (https://ggkbase.berkeley.edu).

### Genome binning, filtering and dereplication

Metagenome assemblies were binned using ggbin. The highest quality bins from each metagenome were selected using DasTool ^75^. Bin quality was manually assessed and refined using the ggkbase platform based on contig coverage, GC values and the inventory of 51 bacterial single-copy genes. Bin completeness and contamination was also analysed using CheckM ^76^ lineage_wf using a threshold of > 70% completeness and < 10% contamination. Finally, bins were dereplicated at 98% nucleotide identity using dRep ^77^.

### Ribosomal protein S3 clustering and diversity analysis

RPxSuite was used as described in Olm et al ^78^ to identify and cluster the universal single-copy ribosomal protein S3 (*rpS3*) gene. The *rpS3* centroid sequences were classified against the GTDB-tk amino acid taxonomic classification database using MMseqs2 ^79^ using the following parameters: easy-taxonomy and --lca-mode 4. *Rps3* was mapped against the reads using BBmap (sourceforge.net/projects/bbmap/) to determine abundance across samples. CoverM ^80^ with a minimum read percent identity of 99% was used to generate a counts table using TPM and normalised by read length and sample.

### Genome phylogenetic classification

GToTree v1.8.7 ^81^ was used to identify, align, trim and concatenate 16 universal ribosomal proteins, defined in Hug, et al ^82^, from the genome set. A maximum likelihood phylogenetic reconstruction of these alignments was generated using IQ-TREE ^83^. The phylogenetic tree was then manually decorated in iTol v6 ^84^.

### PQQ eight-bladed β-propeller identification and classification

For PQQ eight-bladed β-propeller identification and classification of clades, we constructed a phylogenetic tree to discriminate homologous proteins. The PQQ-DH sequences were identified in the dereplicated genome set using a custom HMM for PQQ-binding dehydrogenases as described in Voutsinos et al ^56^. A multiple alignment was generated using MAFFT and the parameters: --local pair --maxiterate 1000 and --reorder. The resulting alignment was trimmed using trimAl ^85^ and the parameter:-gt 0.1. A phylogenetic tree was constructed using IQ-TREE ^83^ and the best-fit model: LG+F+G4. PQQ eight-bladed B-proppeller sequences were manually classified based on their clustering with experimental PQQ dehydrogenase reference sequences and curated sequences taken from Keltjens et al ^20^ and Voutsinos et al ^56^.

### PQQ six-bladed β-propeller identification and classification

The AA12.hmm retrieved from the KEGG KOFAM database was used to identify AA12 (Auxiliary Activity Family 12) PQQ six bladed beta propeller homologs. Retrieved sequences were aligned with the sequence of CcPDH (PDB: 6JWF) and manually searched for all nine PQQ binding residues; R_273_,H_363_,R_430_,N_431_,H_539_,H_560_,W_563_,N_564_,R_621_. InterProScan (v5.51-85.0) was used to search protein sequences against the GENE3D database for the identification of β-propeller domains. AA12 domains were also confirmed by searching sequences against the dbCAN3 database. Sequences were scanned for signal peptides using signalP 6.0. Representative protein sequences were folded using Chai-1 ^86^ using templates and MMseqs2 MSAs. The rank 1 models were profiled against the Protein Data Bank (http://www.rcsb.org/pdb/) (v5e639, date: 2024-04-25) using foldseek ^87^ (v53465) and the easy-search module. The PDB best hits were used for superimposition and residue geometry analyses in ChimeraX (v1.9). Structures were superimposed using the Needleman-Wunsch algorithm and the BLOSUM-62 substitution matrix.

### Protein, protein-protein, and protein-ligand modeling

Proteins were modelled using Chai-1 webserver ^86^ using templates and MMseqs2 MSAs. SMILES text-based notation of PQQ and La^3+^ input as C1=C(C2=C(C(=O)C(=O)C3=C2NC(=C3)C(=O)O)N=C1C(=O)O)C(=O)O and [La+3] were used. The AlphaFold 3 inference pipeline (https://github.com/google-deepmind/alphafold3) was used when modelling large numbers of proteins with ligands in SMILES format. Chai1 and AlphaFold 3 model confidence were evaluated via pLDDT in ChimeraX using the following command: color bfactor palette alphafold. Protein-protein interaction confidence was evaluated in ChimeraX using the following command: alphafold contacts /A to /B. And depth of interaction was measured in ChimeraX using: measure buried area /A with /B. In some cases the signal peptide was identified using SignalP6.0 and excised prior to modelling.

### Protein functional prediction analysis

The tool run_dbCAN v5.2.1 (https://github.com/bcb-unl/run_dbcan) with the parameter ‘easy_CGC’ in protein mode was used to identify CAZyme gene clusters (CGCs) across a set of 3,700 genomes. The CGCs were manually profiled for the presence of PQQ-6β. The ‘substrate_prediction’ function was used to compare identified PQQ-6β containing CGCs against a reference database of characterised polysaccharide utilisation loci (PUL). Finally, gggenes was used to visualise the sequence similarity and domain composition between PQQ-6β CGCs and their PUL best hit as identified by run_dbCAN.

### TBDT, PqqU and PQQ biosynthesis analysis

More than 7000 TonB dependent transporters (TBDTs) were isolated from 704 genomes using a hmmsearch for TBDT pfam PF00593.hmm via hmmer v3.3.1 (https://github.com/EddyRivasLab/hmmer). The identified TBDTs were clustered with five reference TBDT shown to take up REE (Uniport IDs: WP_060849375.1, WP_235726409.1, WP_244426899.1, WP_245268181.1, WP_003609830.1) and *mluA*, the TBDT that is expected to take up methylolanthanin ^44^. Sequences were clustered based on BLASTp all-against-all searches using CLANS ^88^ with an E-value cutoff of 1×10^−10^ until equilibrium. PqqU (PDB ID: 9C40) and PQQ biosynthesis genes were identified following the methods used in Munder, et al ^43^.

### Protein expression, bacterial strain and growth conditions

For plasmid construction, signal peptides were predicted using SignalP 6.0 ^89^ and replaced with a PelB, 10xHisTag and TEV protease and inserted into a pET-28b(+) plasmid. Plasmids were constructed using Twist BioScience. For expression, *E. coli* C41(DE3) cells were transformed with pET-28b(+). A starter culture was inoculated with 90% of the transformation and grown in LB [kanamycin (50 ug/ml)] overnight at 37 °C with rotary shaking at 180 rpm. A contamination control plate was prepared using the remaining 10%. Four liters of TB broth [tryptone (12 g/liter), yeast extract (24 g/liter), K_2_HPO_4_ (12.54 g/liter), KH_2_PO_4_ (2.31 g/liter)], and kanamycin (50 ug/ml) were inoculated with the transformed culture and grown until OD_600_ 1.0, then 0.3 mM IPTG was added to induce expression and incubated overnight at 25 °C in a rotary shaker at 150 RPM.

### Protein purification

Cells were harvested via centrifugation and pellets were resuspended in a final volume of 100 ml of lysis buffer [50 mM Tris-HCl (pH 8), 150 mM NaCl, 20 mM imidazole, lysozyme (0.1 mg/ml), deoxyribonuclease (0.05 mg/ml), and 1× protease inhibitor] and homogenised via pipetting before being subjected to cell lysis in a high-pressure cell disruptor. The lysate was centrifuged at 18,000g for 20 min at 4°C, and the resulting supernatant was loaded on a HisTrap HP 5-ml affinity column. After extensive washing with wash buffer [50 mM Tris-HCl (pH 8), 150 mM NaCl, 20 mM imidazole], the protein was eluted with a step gradient in elution buffer [50 mM Tris, (pH 8), 150 mM NaCl, 500 mM imidazole]. Protein B2 and C5 eluted in 500 mM imidazole. Elution fractions were analysed via SDS-PAGE and pooled and concentrated to a volume of 1 mL in a Amicon Ultra-15 Centrifugal Filter 30-kDa MWCO Millipore. Size exclusion chromatography was performed with a Superdex HiLoad 200 pg 26/600 in size exclusion buffer [50 mM Tris-HCl (pH 8), 150 mM NaCl].

### Protein reconstitution with PQQ, La, and Ca

A 5 mL HiTrap desalting column (Cytiva) was used to buffer exchange proteins B2 and C5 into 50 mM HEPES buffer (pH 7.5) in preparation for ligand incubations. The HiTrap column was flushed with five times column volume of 50 mM HEPES before loading protein. The column was loaded with 1.5 mL purified apoenzyme (8 mg/ml) 50 mM HEPES buffer was injected at a rate of 5 ml/min and the first 2 mL fractions were collected. The apoenzymes in 50 mM HEPES were incubated with a 5-fold molar excess of methoxatin (PQQ), LaCl_3_ and CaCl_2_ for 30 minutes at room temperature. Incubated samples were then passed through the HiTrap column again to remove unbound PQQ, La and Ca. Protein concentration was measured after each column passthrough using the NanoDrop Lite Plus (thermo scientific).

### Identification of protein PQQ and metal content

The absorbance spectrum of the reconstituted enzyme was recorded in a 1-cm-path-length quartz cuvette at room temperature using a Cary 60 UV-Vis spectrophotometer (Agilent Technologies). Five technical repeats were measured for each protein using inductively coupled plasma mass spectrometry (ICP-MS) on a Agilent LC/ 8900 ICP-QQQ mass spectrometer at Bio21, Melbourne University. Lanthanum concentrations were quantified using 0-500 ppb lanthanum standards.

## Data availability

All Supplementary Data Files and sequencing data will be made available upon publication.

## Author contributions

M.Y.V., J.F.B., and R.G. conceived and designed the experiments. M.Y.V. performed the experiments. M.Y.V, J.F.B, R.G., and C.M.R. analysed the data. M.Y.V. and J.F.B. wrote the paper. All authors reviewed the final manuscript.

## Supplementary Figures

**Figure S1.**
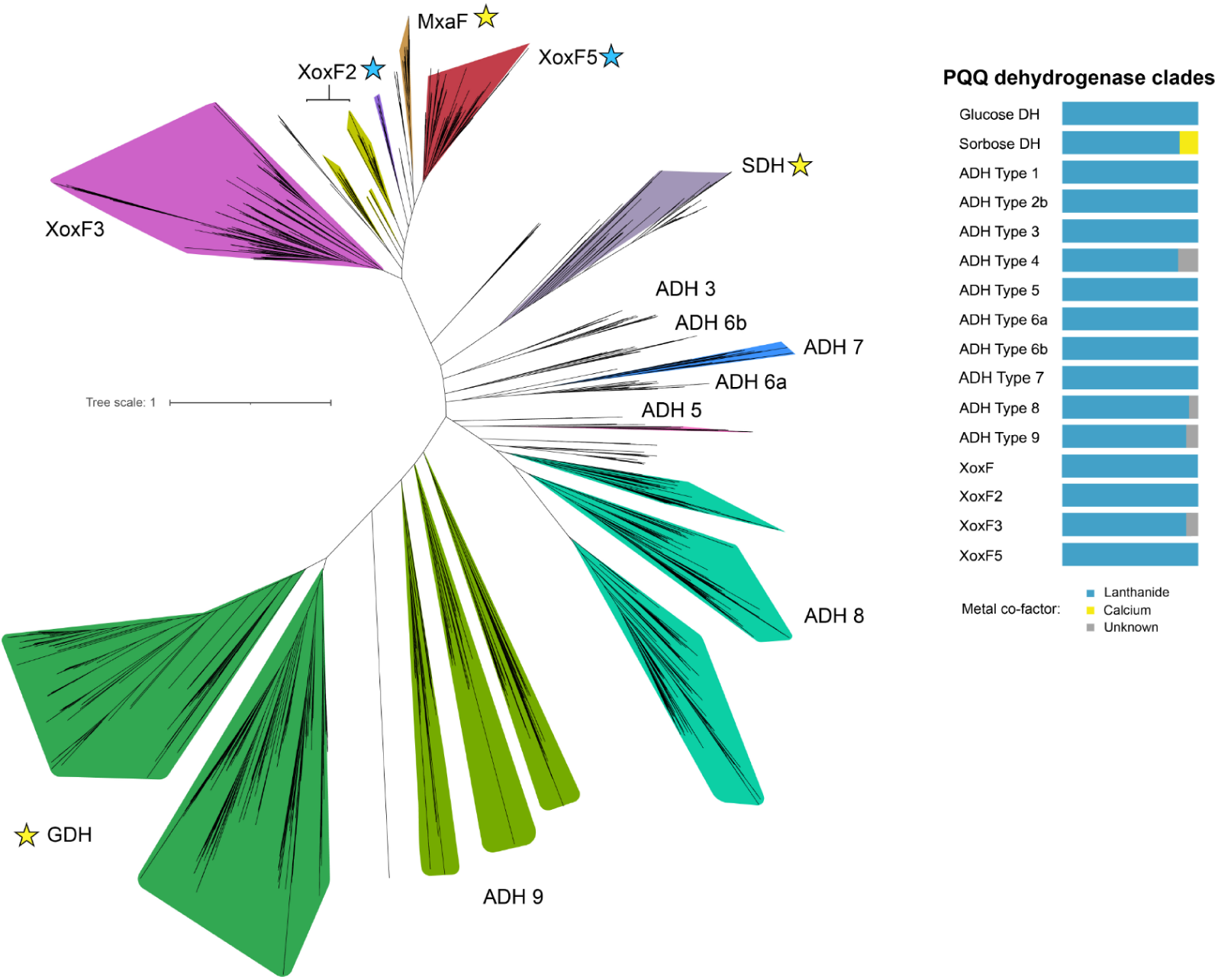
Unrooted phylogenetic tree of PQQ eight-bladed β-propeller proteins identified in genomes reconstructed from weathered granitic rock. Stars indicate if a group contains an experimentally confirmed protein, yellow = Ca dependent, blue = Ln dependent. SDH = sorbose dehydrogenase, ADH = alcohol dehydrogenase, GDH = glucose dehydrogenase. Legend shows the ratio of proteins in each group that bind Ln = blue, yellow = Ca, or grey = unknown.

**Figure S2.** Rectangular phylogenetic tree of PQQ eight-bladed β-propeller proteins identified in genomes reconstructed from weathered granitic rock. Sequences from this study begin with ‘RPR’ in the label. The active site aspartate residues (DxD) from each sequence are displayed. This figure is available via figshare at: 10.6084/m9.figshare.32955410.

**Figure S3.** Rectangular view of phylogenomic tree from. Fig 1. This figure is available via figshare at: 10.6084/m9.figshare.32955410.

**Figure S4.**
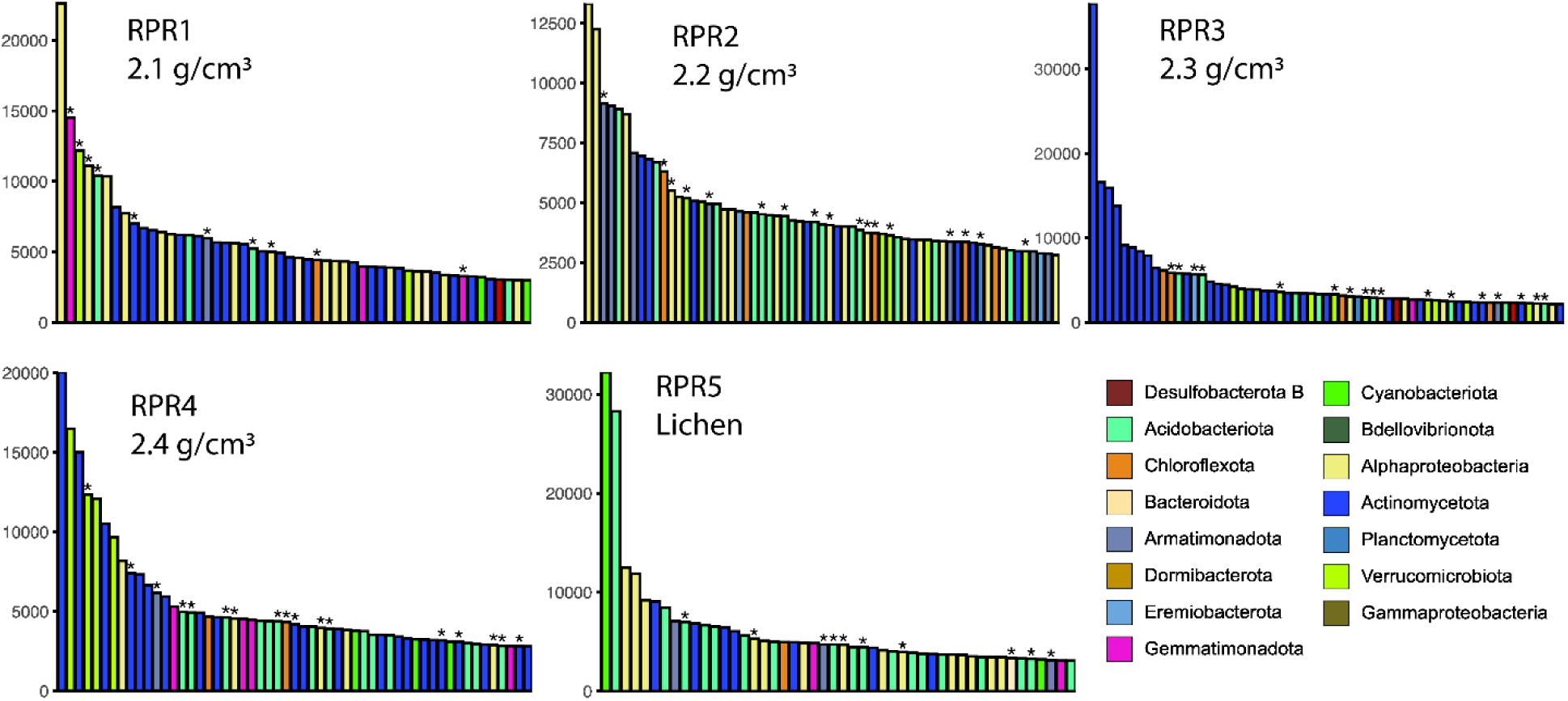
Rank abundance curves of the top 30% most abundant species (rpS3) across each sample. Read counts are expressed as TPM. Asterisks above bars indicate binned rpS3 that contain Ln-dependent PQQ enzymes.

**Figure S5.**
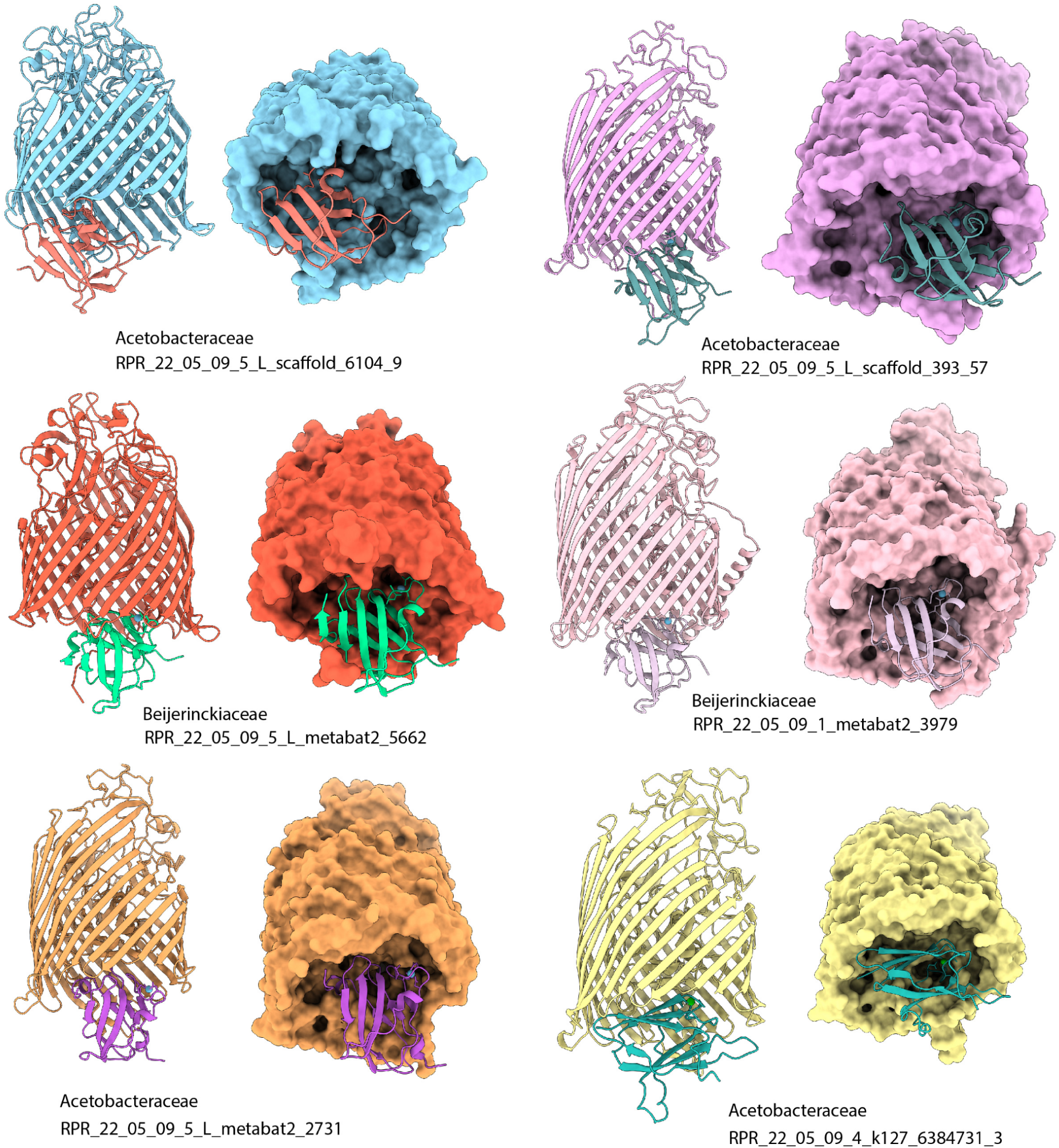
**The predicted interaction between putative lanthanide periplasmic binding proteins and their associated TonB transporter.**

**Figure S6.**
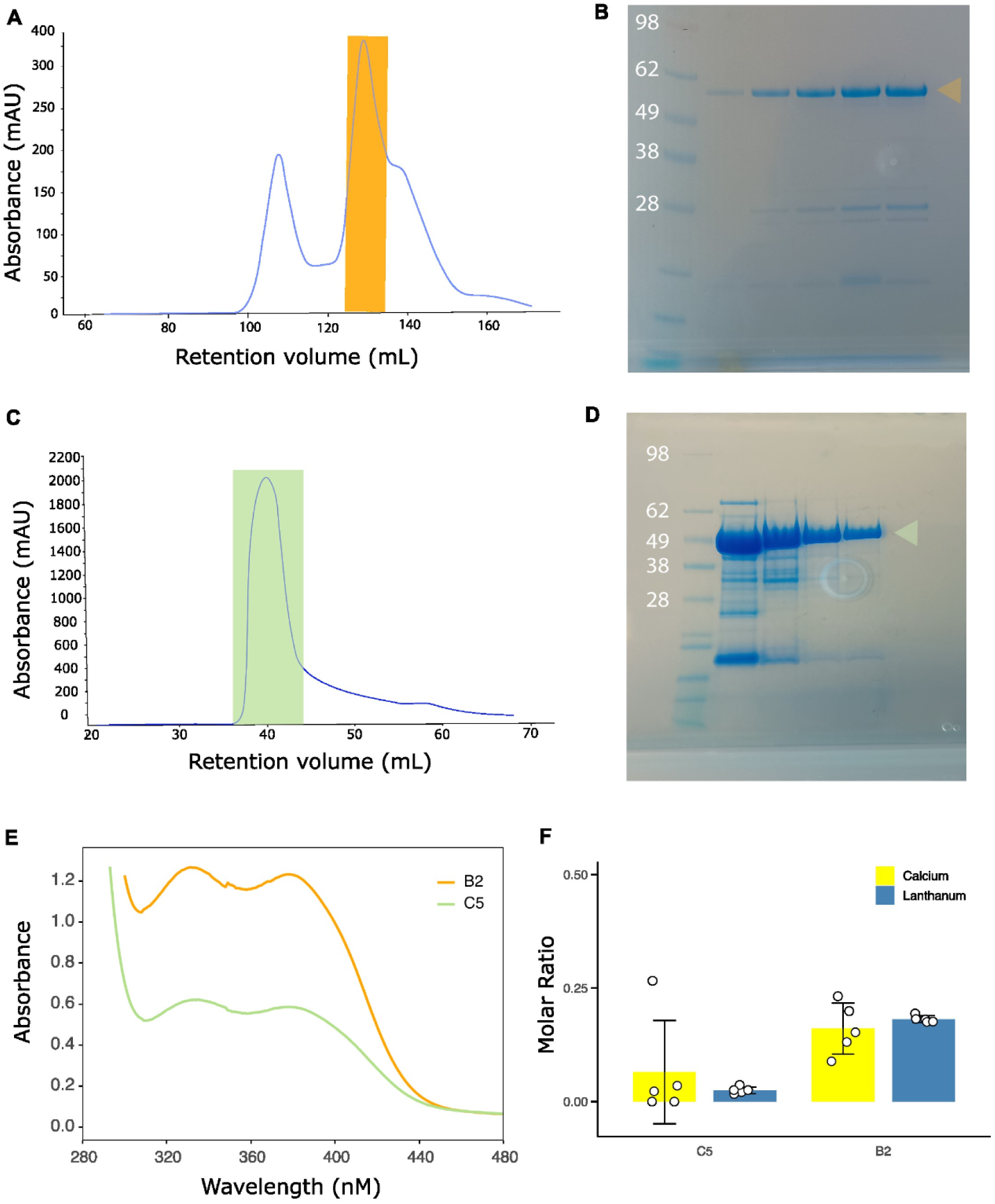
A) Size exclusion chromatography elution profile of RPR_22_05_09_4_k127_6634144_13 (B2). B) SDS-PAGE gel of purified B2. C) Size exclusion chromatography elution profile of RPR_22_05_09_1_scaffold_1078_6 (C5). D) SDS-PAGE gel of purified RPR_22_05_09_1_scaffold_1078_6 (C5) E) UV-VIS spectrums of C5 and B2 after incubation with PQQ. F) ICP-MS results from untreated C5 and B2 controls. Error bars, standard deviation. White circles, molar ratio of five technical replicates. The height of the bar graph is the mean of molar ratios.

**Fig S7.**
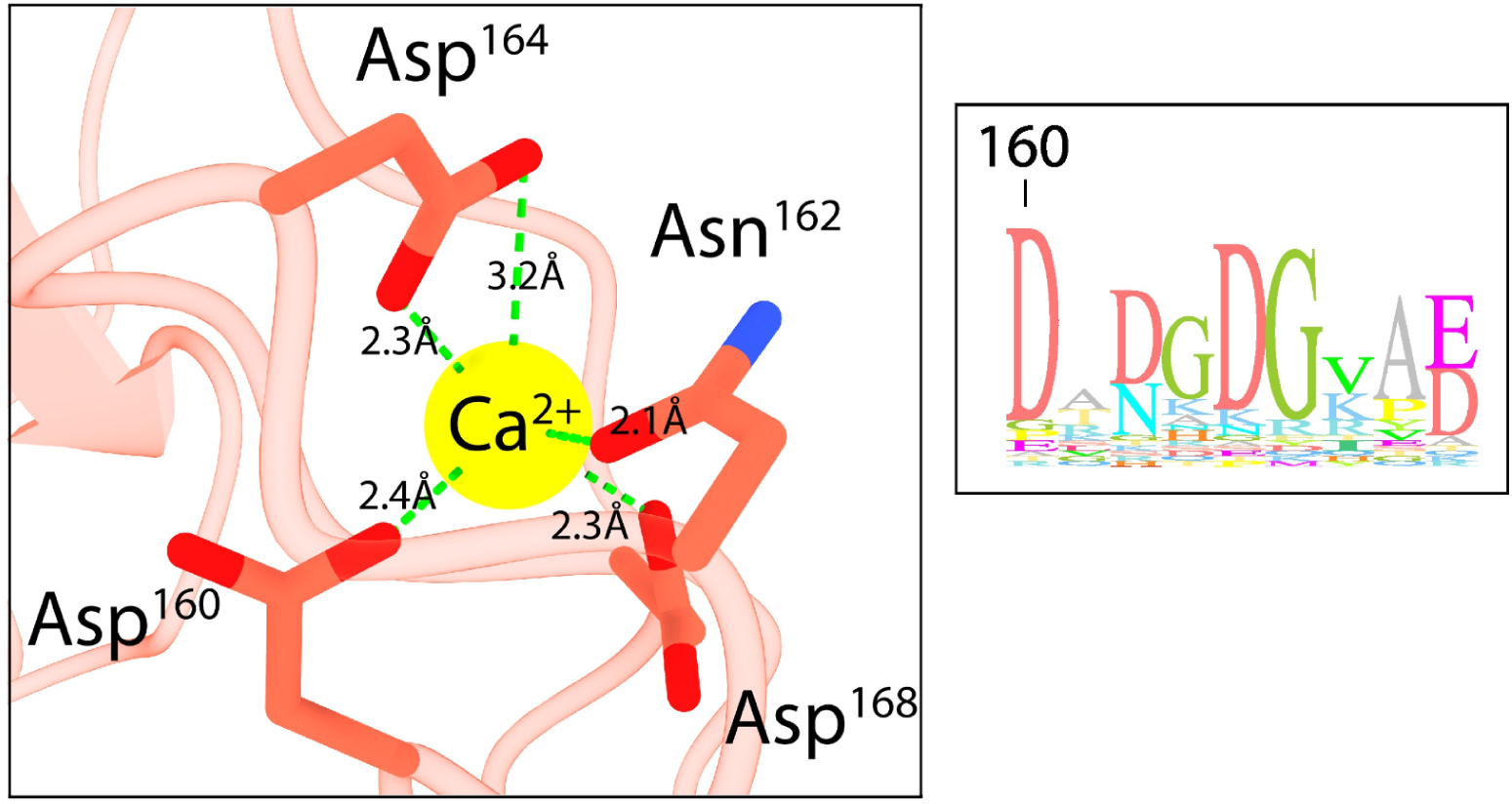
The predicted second metal binding site of B2. The sequence logo insert illustrates the frequency of the predicted metal binding residues across 21,993 PQQ-6β sequences.

**Figure S8.**
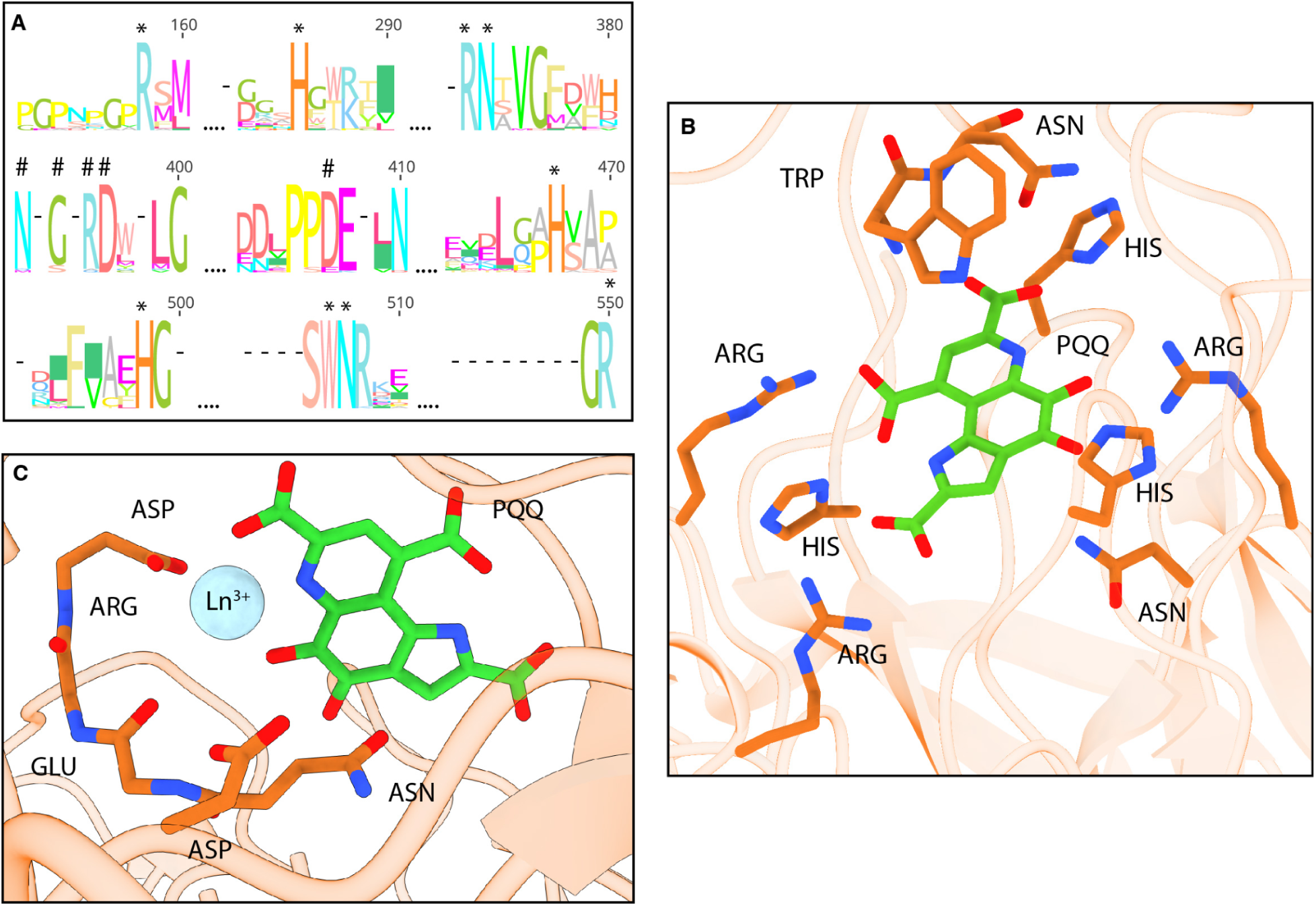
Predicted lanthanide binding in archaeal PQQ-6β-propellers. A) Consensus sequence logo of 158 archaeal sequences highlighting the conserved residues responsible for binding PQQ (‘*’) and a metal cofactor (‘#’). B) Predicted residues involved in PQQ binding in an archaeal homolog (CABMIA010000186.1_15). C) Zoomed-in view of the metal binding active site illustrating the residues involved in lanthanum coordination. Only the backbone carbonyl oxygen of Glu and Arg is shown and predicted to be involved in metal coordination.

**Figure S9.** PQQ-6β proteins in CAZyme gene clusters from bacterial genomes. This figure is available via figshare at: 10.6084/m9.figshare.32955410.

## Supplementary tables

**Table S1.** Whole-rock elemental composition across a weathered granite profile. Values are in PPM. S# = sample number. For full whole-rock elemental composition see Data S1.

|  | RPR8(S5) | RPR4 (S4) | RPR3(S3) | RPR2(S2) | RPR1(S1) |
| --- | --- | --- | --- | --- | --- |
| Calcium | 18,023 | 17,798 | 22,426 | 16,809 | 15,607 |
| Iron | 31,182 | 26,926 | 29,392 | 30,782 | 25,952 |
| Phosphate | 548 | 241 | 231 | 1 | 1 |
| Lanthanum | 15 | 45 | 64 | 72 | 82 |
| Cerium | 64 | 171 | 130 | 132 | 147 |
| Neodymium | 20 | 40 | 55 | 58 | 66 |
| Titanium | 1 | 1 | 1 | 1 | 1 |

**Table S2.** Distance of active site residues to metal ion (Å). * Coordinates with metal via a water molecule. ^-^ Does not coordinate with metal.

| Atom | Distance to La (B2) | Atom | Distance to La (C5) | Atom | Distance to Ca (CcPDH, PDB: 6JWF) |
| --- | --- | --- | --- | --- | --- |
| Asn<sup>307</sup> OD1 | 5.0 | Asn<sup>264</sup> OD1 | 6.5 | *Asn<sup>448</sup> OD1 | 4.7 |
| Gly<sup>308</sup> O | 2.8 | Glu<sup>265</sup> O | 2.6 | Ser<sup>449</sup> O | 2.5 |
| Arg<sup>309</sup> O | 4.2 | Arg<sup>266</sup> O | 4.2 | *Leu<sup>450</sup> O | 4.2 |
| Asp<sup>310</sup> OD1 | 2.3 | Asp<sup>267</sup> OD1 | 2.3 | Asp<sup>451</sup> OD1 | 2.3 |
| Asp<sup>310</sup> OD2 | 3.7 | Asp<sup>267</sup> OD2 | 3.9 | †Asp<sup>451</sup> OD2 | 4.0 |
| Asp<sup>319</sup> OD1 | 4.6 | Asp<sup>276</sup> OD1 | 4.6 | Glu<sup>471</sup> OE1 | 4.9 |
| Asp<sup>319</sup> OD2 | 5.5 | Asp<sup>276</sup> OD2 | 5.3 | *Glu<sup>471</sup> OE2 | 4.7 |
| PQQ O5 | 2.1 | PQQ O5 | 2.5 | PQQ O5 | 2.4 |
| PQQ N6 | 2.4 | PQQ N6 | 2.4 | PQQ N6 | 2.5 |
| PQQ O7 | 2.4 | PQQ O7 | 2.3 | PQQ O7 | 2.4 |

